# Beyond rhythm – A framework for understanding the frequency spectrum of neural activity

**DOI:** 10.1101/2023.05.12.540559

**Authors:** Quentin Perrenoud, Jessica A. Cardin

## Abstract

Cognitive and behavioral processes are often accompanied by changes within well-defined frequency bands of the local field potential (LFP i.e., the voltage induced by neuronal activity). These changes are detectable in the frequency domain using the Fourier transform and are often interpreted as neuronal oscillations. However, aside some well-known exceptions, the processes underlying such changes are difficult to track in time, making their oscillatory nature hard to verify. In addition, many non-periodic neural processes can also have spectra that emphasize specific frequencies. Thus, the notion that spectral changes reflect oscillations is likely too restrictive. In this study, we propose a simple yet versatile framework to understand the frequency spectra of neural recordings. Using simulations, we derive the Fourier spectra of periodic, quasi-periodic and non-periodic neural processes having diverse waveforms, illustrating how these attributes shape their spectral signatures. We then show how neural processes sum their energy in the local field potential in simulated and real-world recording scenarios. We find that the spectral power of neural processes is essentially determined by two aspects: 1) the distribution of neural events in time and 2) the waveform of the voltage induced by single neural events. Taken together, this work guides the interpretation of the Fourier spectrum of neural recordings and indicates that power increases in specific frequency bands do not necessarily reflect periodic neural activity.

## Introduction

The Fourier transform, or analogous methods, is routinely used to analyze the frequency content (i.e., the spectrum) of neural activity (Penttonen and Buzsáki, 2003; Bruns, 2004). Frequency spectra are highly sensitive to changes in the processes driving a signal (Percival and Walden, 1993). They can thus be used to detect subtle variations in the dynamics of the brain’s electric fields during attention, memory formation or retrieval, locomotion, and motor responses (Gray *et al*., 1989; O’Keefe and Recce, 1993; Skaggs *et al*., 1996; Fries *et al*., 2001; Girardeau *et al*., 2009; Niell and Stryker, 2010; Bosman *et al*., 2012; Vinck *et al*., 2015; Chen *et al*., 2017; Veit *et al*., 2017; Uran *et al*., 2022). However, the behavior of frequency spectra is often counterintuitive. For instance, the finite nature of recordings induces distortion of their spectral content (Percival and Walden, 1993). Furthermore, relating frequency spectra to continuously evolving neural processes is a complex problem. As a result, the interpretation of the frequency spectrum of neural activity is not straightforward.

Increased spectral power within some frequency bands is often believed to reflect oscillations. In fact, the words “oscillation” and “rhythm” are generally used to describe the subfield of neuroscience dedicated to study of the brain’s electrical activity (Buzsáki and Vöröslakos, 2023). Some neural patterns do indeed show strong periodicity. This includes well-characterized rhythms such as hippocampal theta (Vanderwolf, 1969; Winson, 1974; O’Keefe and Recce, 1993; Skaggs *et al*., 1996), thalamic spindles (Steriade, 2006; Niethard *et al*., 2018) and slow waves observed in the cortex during sleep (Steriade, Nuñez and Amzica, 1993; Contreras, Timofeev and Steriade, 1996; Sanchez-Vives and McCormick, 2000). Nonetheless, neuronal processes can also display non-periodic dynamics (Burns, Xing and Shapley, 2011; Xing *et al*., 2012; Weber and Pillow, 2017; Naud and Sprekeler, 2018; Donoghue *et al*., 2020; Williams *et al*., 2021; Spyropoulos *et al*., 2022). In addition, most signals, periodic or not, have spectra where some frequencies are enhanced. Gaussian functions are non-periodic, yet exhibit a spectrum with a strong representation of low frequencies (Starosielec and Hägele, 2014). Thus, the notion that oscillations underlie changes in the spectrum of neural activity appears generally restrictive.

Here, we provide a broadly applicable conceptual framework and a didactic discussion for the often-intimidating interpretation of the frequency spectrum of neural recordings. We show that neural patterns are usefully conceptualized as processes where discrete events can occur with varying degrees of periodicity. Using simulations, we illustrate how multiple processes can sum up in the Local Field Potential (LFP) and how the basic properties of the Fourier transform shape the frequency spectrum of neural recordings. We show that the spectral signature of neural patterns depends essentially on 2 aspects: 1) the distribution of events in time and 2) the waveform of the voltage induced by individual events. Finally, we show examples of how this conceptual framework can be applied to decipher processes inducing changes in the spectral profile of real-world data acquired in the primary visual cortex of behaving mice.

## Materials and Methods

All simulations and analysis were performed in Matlab 2018b (Mathworks). Simulated time series had a sampling rate of 1kHz and a duration of 1000 seconds. Event timing pulse trains had a value of 1 at the time of events and zero everywhere else. Except in the case of perfectly periodic processes, event intervals were drawn from a gamma distribution:

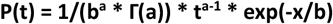

where **P(t)** is the probability density of the interval **t, Γ(x)** is the gamma function of the input value **x** (i.e., the gamma function is a generalization of the factorial to real numbers), and **a** and **b** are parameters determining the shape of the distribution. The process is super-Poissonian If **a** < 1, Poisson if **a** = 1 and super-Poissionian if **a** > 1. Parameter **b** was always set to **1** / (**a** ^**\***^ **f**) where **f** is the overall event frequency of the process. Values of **a** and **f** were as follows: Figure 1A: **a** = 0.1, **f** = 40Hz; Figure 1B: **a** = 1; **f** = 40Hz; Figure 1C; **a** = 31.6228; f = 40Hz; Supplementary Figure 1A: **a** = 1, **f** = 40Hz; Supplementary Figure 1B: **a** = 10, **f** = 40Hz; Supplementary Figure 1C: **a** = 100, **f** = 40Hz; Supplementary Figure 1D: **a** = 1000, **f** = 40Hz; Supplementary Figure 2A: **a** = 1, **f** = 50Hz; Supplementary Figure 2B: **a** = 20, **f** = 80Hz; Supplementary Figure 2C: **a** = 40, **f** = 20Hz; Supplementary Figure 4A: **a** = 15.8489, **f** = 40Hz; Figure 3B: **a** = 1, **f** = 75Hz.

**Figure 1:**
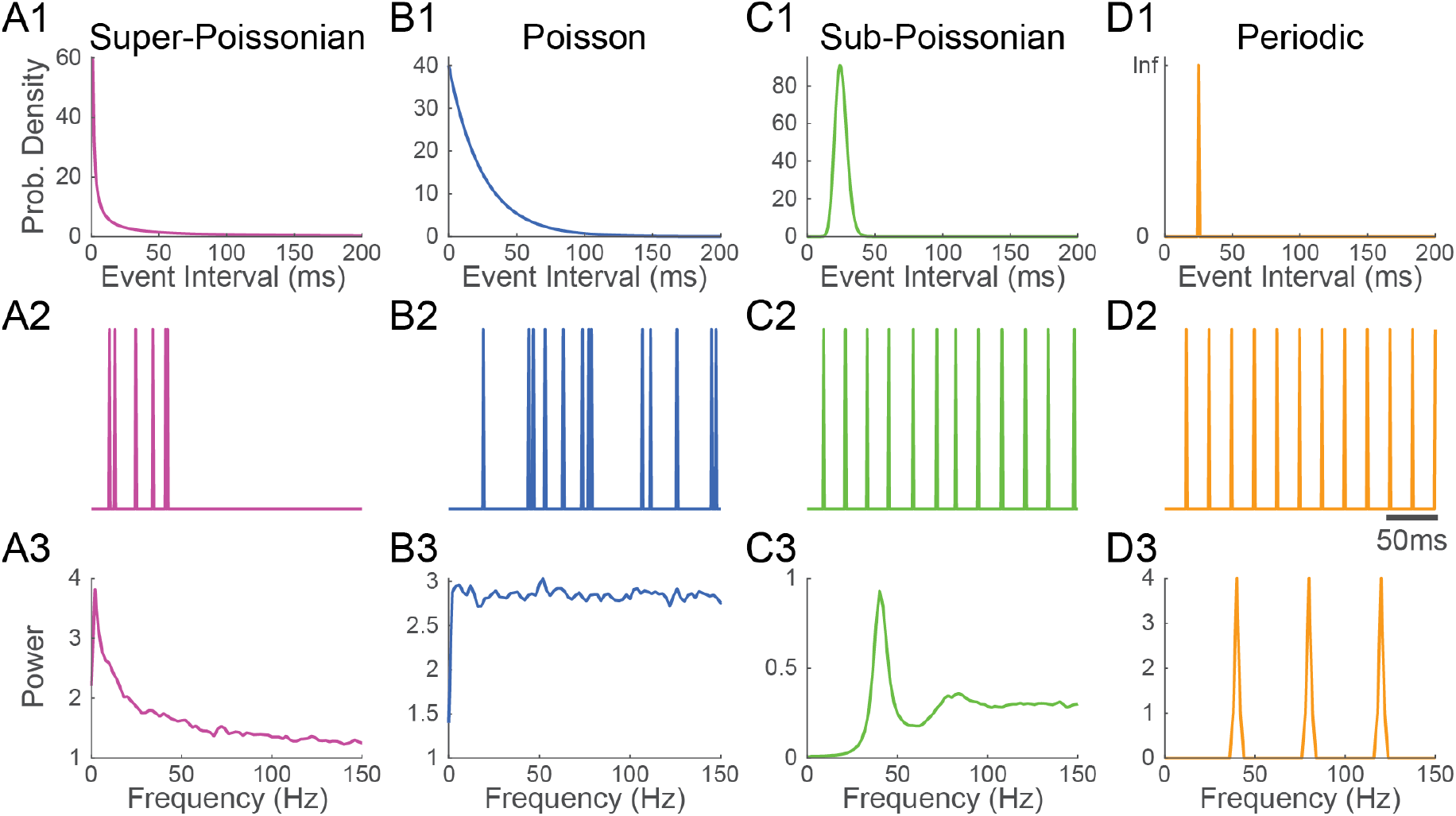
Super-Poissonian, Poisson, sub-Poissonian and periodic point-processes have recognizable Fourier spectra. **1)** Event interval distribution, **2)** excerpt of an event train and **3)** Fourier spectrum of **A)** super-Poissonian, **B)** Poisson **C)** sub-Poissonian and **D)** perfectly periodic point processes. Distinct interevent interval distributions translate into specific energy distribution in the Fourier spectrum. All processes have a mean frequency of 40Hz.

Recurring neural events were simulated by convoluting event pulse trains with waveform traces of 500ms. Various waveforms (i.e., impulse response functions) were considered. Synaptic events were modelled with the alpha function (Markram *et al*., 1997):

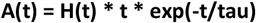

Where **A(t)** is the alpha function at time **t, H(t)** is Heaviside step function, and **tau** is a parameter determining the time course of the synaptic response. Parameter **tau** was always set to 5.6ms.

Spikes and spindles were modelled with real valued Morlet wavelets (i.e. the multiplication of a sinusoid with a gaussian window):

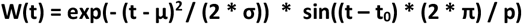

Where **W(t)** is the Morlet wavelet at time **t, μ** and **σ** are the mean and standard deviation of the gaussian component, **t**_**0**_ is the offset of the sine function and **p** is its period. Parameter **μ** was always set to zero (i.e., centered). Values of **σ, t**_**0**_, and **p** were as follows: Figure 2C: **σ** = 6.25ms, **t**_**0**_ = 13.75ms, **p** = 25ms; Figure 2D: **σ** = 25ms, **t**_**0**_ = 0ms, **p** = 25ms; Supplementary Figure 3: same as Figure 2; Figure 3B: **σ** = 3.33ms, **t**_**0**_ = 7.33ms, **p** = 13.33ms.

**Figure 2:**
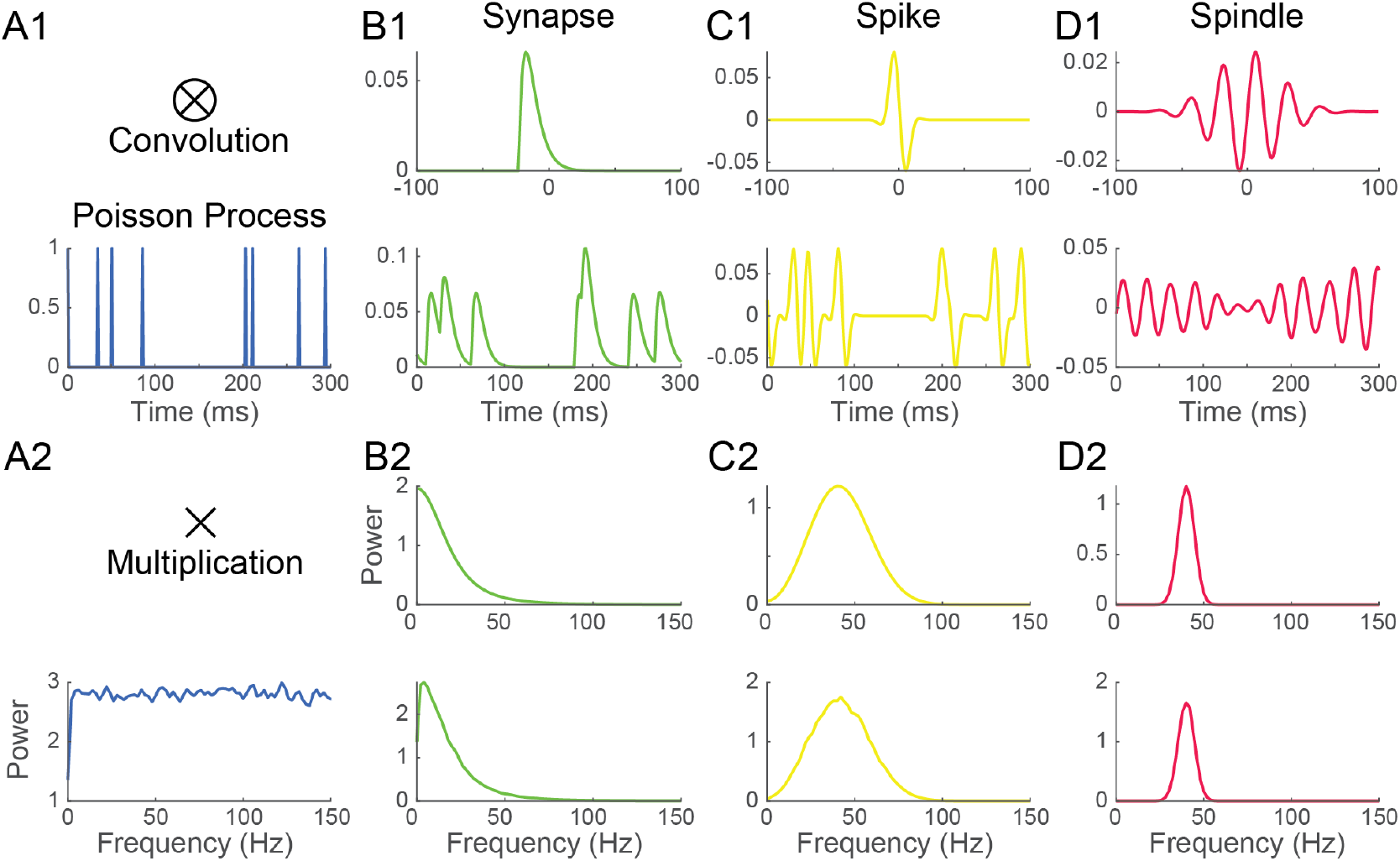
The Fourier spectrum of recurring event trains depends on the inter-event interval distribution and the waveform of single events. **1) A)** a Poisson process was convolved with 3 waveforms (top) mimicking the shape of **B)** a synaptic event, **C)** a spike and **D)** a spindle, resulting in three distinct recurring event time series having the same event timing (bottom). **2)** Convolution in the time domain translates into a simple multiplication into the frequency domain. Thus, the Fourier spectrum of the recurring event time series in B1, C1 and D1 (bottom) is simply the product of the spectrum of their waveform (top) and the spectrum of the Poisson pulse train (A2). Here, as Poisson processes have a flat spectrum, the spectrum of event times series is essentially determined by that of their waveform.

**Figure 3:**
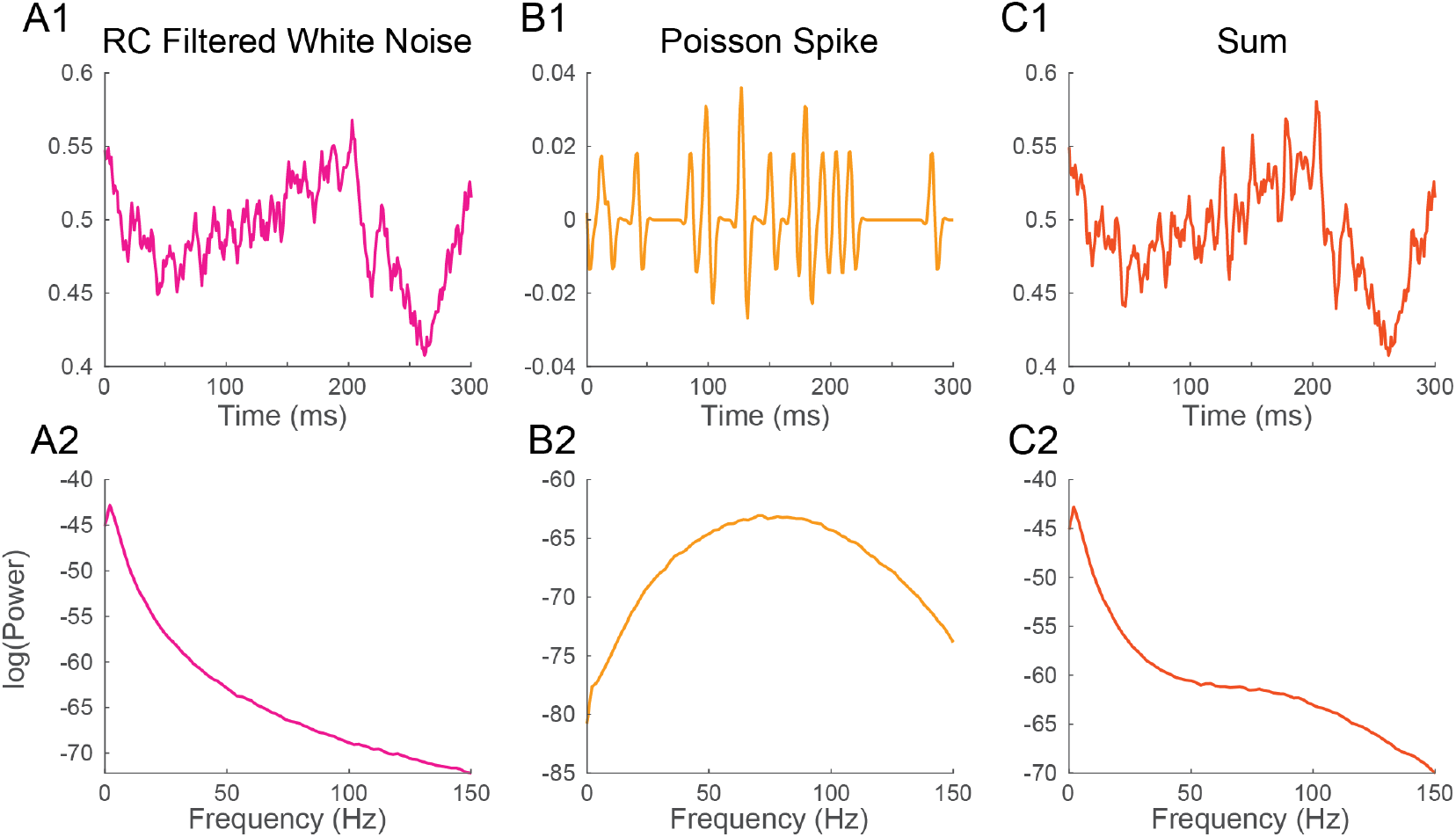
Neural processes sum their energy in the Fourier spectrum. **1)** Excerpt, and **2)** Fourier spectrum of **A)** white noise filtered with a passive Resistive-Capacitive (RC) filter mimicking desynchronized background synaptic activity, **B)** a Poisson spike train and **C)** the sum of the time series in A and B. The spectrum of background synaptic activity in A has a characteristic 1/f^2^ shape which is inherited from the RC filter. The spectrum of the Poisson spike train in B has a bell shape peaking at 75Hz which is inherited from the spike waveform. The spectrum of the sum of two signals is the sum of their spectra.

To study the effect of waveform width, waveforms were modelled as Hann window function (i.e. one cycle of a sinusoid) with periods of 5ms, 25ms and 125ms (Supplementary Figure 4B, 4C and 4D, respectively).

Background synaptic noise was modelled by convolving white noise with a 500ms waveform corresponding to the impulse response function of a passive resistive capacitive (RC):

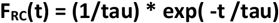

Where **F**_**RC**_**(t)** is the value of the impulse response function of the RC filter at time **t**, and **tau** is a parameter determining the time course of the response (note the similarity to the alpha synaptic response). Parameter **tau** was set to 35ms based on values inferred from recording of the membrane resistance and capacitance of excitatory neurons in the rat barrel cortex (Karagiannis *et al*., 2009).

The spectrum of waveforms was computed by tapering with a Hann window and taking the squared amplitude of the Fourier transform. The spectrum of recurring event time series was estimated with Welch’s method, that is by averaging the squared amplitude of the Fourier transform within non-overlapping 500ms segments tapered with a Hann window.

The experiment described in Figure 4 is part of a data set used in (Perrenoud *et al*., 2022). The method used for the extraction of gamma events is detailed and freely available at (https://github.com/cardin-higley-lab/CBASS).

**Figure 4:**
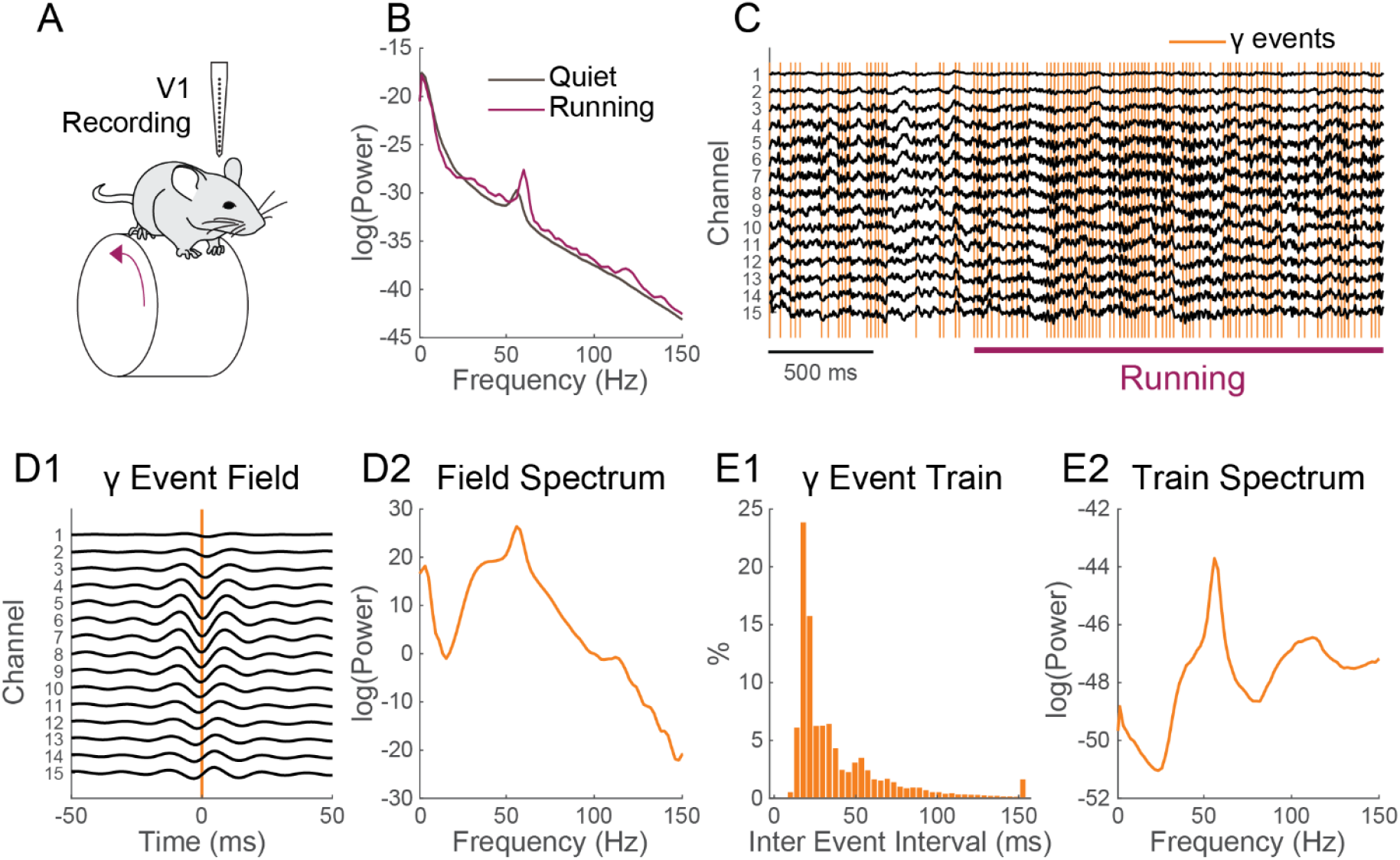
The spectrum of gamma activity in the mouse visual cortex is shaped by the timing of gamma events. **A)** Schematic of the recording configuration. A linear electrode array is used to record the Local Field Potential (LFP) across the layers of the primary visual cortex (V1) of a mouse freely running on a wheel. **B)** Locomotion (purple) induces an increase in LFP power in the gamma range (30-80Hz) having a narrow band peak around 55-60Hz and broad band component. **C)** Excerpt of the recording around locomotion onset. Locomotion modulated gamma activity can be tied to discrete network event (orange; further detail on the method can be found in Perrenoud et al. 2022). **D) 1)** Average LFP around gamma events and **2)** its Fourier spectrum. The average LFP shows a broad band and a narrow band peak in its energy distribution. **E) 1)** Inter-event interval distribution of gamma events and **2)** Fourier spectrum of the event timing train of gamma events. The broad-band and a narrow-band components of locomotion modulated gamma activity are visible in the Spectrum of the event’s timing.

## Results

To understand the frequency spectrum of neural recordings, it is useful to first consider the origin of the brain’s electric field. Neurons encode signals through variations of their membrane potential which are induced by the diffusion of Na+, K+, Cl− and Ca2+ ions through tightly controlled ion channels (Buzsáki, Anastassiou and Koch, 2012). This movement causes variations in the electric field whose behavior is described by Maxwell’s equations. With a perfect knowledge of the brain’s electric (and magnetic) field, Maxwell’s equations might in principle, permit to infer movements of charges at the neuronal level. However, the sensitivity of our recording methods and the complexity of ion movement through the several billions of neurons that constitute the brain make this intractable in practice.

While the brain’s electric activity can give the impression of a continuously evolving chaos, it is made out of many repeating elements. These elements are the activation of synapses, the firing of neurons or, at a larger scale, coordinated firing and synaptic barrages within one brain region, or from one area to another (Womelsdorf *et al*., 2014). Each such event has a defined impact on the brain’s electric field and may thus induce a voltage deflection in a recording electrode. A recurring event, such as the firing of one neuron, can at time have of a strong influence on a signal depending on its magnitude and its proximity (Schmitzer-Torbert *et al*., 2005). We will thus start by considering how one recurring event affects the spectrum.

### Impact of a recurring event’s temporal distribution on the Fourier spectrum

Let us first examine the influence on the spectrum of how often and regularly an event occurs. To assess this, we will idealize recurring events and treat them as point processes. Point processes are completely defined by their inter-event interval distribution. The simplest and most commonly occurring type of point-process is the Poisson process. Poisson processes are characterized by a decaying exponential interevent interval distribution (Figure 1B) and have 2 interesting properties: 1) the probability of an event at any given time is invariant and entirely unaffected by the process’s history; 2) the variance of the number of events per unit of time is equal to the event rate.

Point processes can be generally categorized by how regularly events occur compared to a Poisson process. Processes happening more regularly have a less variable inter-event interval. The variance of the number of events per unit of time in thus lower than the event rate. These processes are called sub-Poissonian (Figure 1C). As the variance of the event interval drops, the process begins to resemble a periodic process (Supplementary Figure 1). In the most extreme case, the variance of the inter-event interval is null, and the process is perfectly periodic (Figure 1D). Conversely, the variance of the number of events per unit of time can be superior to the event rate. In this case, the process is called super-Poissonian (Figure 1A). The firing of bursting neurons is an example of super-Poissonian process (Williams *et al*., 2021).

The gamma distribution (among others) can be parameterized to generate point processes (called gamma processes) having Poisson, super-Poissonian or sub-Poissonian inter-event intervals. To illustrate how the variance of point-processes affect their frequency spectrum, we parameterized gamma processes to generate super-Poissonian, Poisson and sub-Poissionian event time series. A perfectly periodic event times series was also constructed. All four time-series had a simulated sampling rate of 1000Hz and a matched event rate of 40Hz (Figure 1). The Fourier spectrum of each time series was then estimated using Welch’s methods with a 500ms Hann window (Materials and Methods), a method classically used to estimate Fourier spectra in real data.

As can be seen in Figure 1B3, the spectrum of a Poisson train of event is perfectly flat. While the rate of a Poisson process affects overall power (energy per unit of time), energy remains evenly distributed across frequencies. The spectrum of a super-Poissonian gamma process is a decaying exponential and has thus more energy at low frequencies (Figure 1A3). Conversely, the spectrum of sub-poissonian gamma process shows reduced power at low frequency, a pronounced peak around the rate of the process and a flat energy distribution for higher frequencies (Figure 1C3). As the process becomes more regular secondary peaks start appearing at the harmonics of the processes rate (i.e., integer multiples of the core frequency, in our case 80Hz, 120Hz and so on; Supplementary figure 1). For a perfectly periodic point process, the power is entirely concentrated and evenly distributed across the process’s rate and its harmonic (Figure 1D3). It thus appears that the event distribution of a process has a substantial and recognizable influence on the shape of its spectrum.

Periodic, Poisson and sub or super-Poissionian Gamma processes are idealized processes that might not be representative of what happens in the brain. To illustrate how the timing of events affect the Fourier spectrum in more arbitrary cases, we built a compound inter-event interval distribution by adding the distributions a Poisson process and 2 sub-Poissonian gamma processes having distinct rate and variance (Supplementary Figure 2). Accordingly, the inter-event interval distribution showed two prominent modes on top of an exponential decay. We generated an event times-series following this distribution and for each summed process for comparison. The Fourier spectrum of each time-series was computed as described above.

The Fourier spectrum of the compound process (Supplementary Figure 2D3) closely resembled the sum of the Fourier spectra of the summed process (Supplementary Figure 2A3, B3, C3). Two main peaks were observed respectively at the frequencies of each sub-Poissonian process. These results illustrate that, for a point process, peaks in the spectrum are broadly indicative of modes (i.e., preferred intervals) in the inter-event interval distribution.

### Impact of a recurring event’s waveform on the Fourier spectrum

So far, we have only considered idealized time series where the impulse response function (i.e., the waveform of the voltage induced by one event) of single events is a perfect pulse (i.e., having a value of 1 at the time of the event and zero everywhere else). In actual recordings of the brain’s field activity, repeating events will have more complex waveforms. We will now consider how different waveforms affects the Fourier spectrum in conjunction with the event interval distribution. For the sake of simplicity, we will only consider that the waveform of all events in a process does not vary in shape or amplitude.

A train of recurring events can be seen as the convolution of its waveform with a signal representing the events timing such as the point-processes considered in the previous section. The Fourier transform has the remarkable property that the spectrum of the convolution of two time series is the product of their spectrum. In other words, convolution in the time domain translate into a simple multiplication in the frequency domain. Within the accuracy of estimation methods, the Fourier spectrum of a time series of recurring events is thus equivalent to the product of the spectra of the event’s waveform and that of its event timing pulse train.

To illustrate how this interaction shapes the Fourier spectrum, we considered three basic waveforms: 1) the alpha function, a popular choice to model synaptic responses (Figure 2B1); and 2 real-valued Gabor wavelet (the product of a sinusoidal and a gaussian window) shaped respectively to resemble 2) a spike (Figure 2C1) and 3) a spindle (Figure 2D1). All impulse response functions have specific Fourier spectra (Figure 2B2, C2 and D2). The alpha function has a spectrum where energy is concentrated in lower frequencies. The spectrum of Gabor wavelets is centered on the frequency of their sinusoid component and has a bandwidth inversely proportional to the width of their gaussian window.

Each impulse response function was then convolved with a Poissonian (Figure 2) or a periodic timing pulse train (Supplementary Figure 3). The Fourier spectrum of resulting time-series was estimated as before. As predicted by theory, all constructed event time-series had spectra nearly equal to the product of that of their timing pulse train and that of their waveform. Due to the flat shape of their timing pulse train’s spectrum, Poisson distributed event time-series have a Fourier spectrum that is essentially that of their waveform (Figure 2B2, C2 and D2). Thus, as exemplified in the Poissonian spikes train constructed in Figure 2C, a pronounced peak in the spectrum can be inherited solely from the shape of a recurring event’s voltage response without periodicity in its timing.

Contrasting with Poisson trains, the energy of periodic timing pulse trains is concentrated at the process’s core frequency and its harmonics. Thus, in this case, the spectrum is more generally determined by the spectrum of the timing pulse train than by that of the waveform. However, as exemplified in Supplementary Figure 3B2, some waveforms will greatly reduce the power of a periodic timing pulse train when the energy distribution of their spectrum does not overlap.

The examples above illustrate that the spectrum of a train of recurring events is shaped both by the event timing (i.e. its timing pulse train) and the impact of single events on the signal (i.e. its impulse response function or waveform). How much each shapes the spectrum varies depending on the case. One general parameter will however influence the relative contribution of one over the other, namely the duration of the waveform relative to an events overall rate of occurrence.

To illustrate this, we considered a moderately periodic sub-Poissonian pulse train (Supplementary Figure 4). The train was then convolved with Hann functions (i.e., one cycle of a sinusoid and a popular window function for spectral estimation) of increasing width relative to the processes core frequency (40Hz). For short Hann pulses (5 times shorter the average inter-event interval, Supplementary Figure 4B), the spectrum of the resulting event times series closely resembles that of the timing pulse train. However, for longer waveforms (5 times longer the average inter-event interval, Supplementary Figure 4D), the shape of the spectrum was mostly determined by that of the waveform.

These results indicate that neural processes with short waveforms, such as action potentials, have a spectrum that tends to be influenced by their timing (i.e. timing pulse train). Conversely, neural process with a longer waveform relative to their rate of occurrence, such as synaptic events, will have a spectral imprint that tends to be determined by the waveform of single events (i.e. impulse response function).

### How do multiple neural processes sum up in the Fourier spectrum?

We have examined the Fourier spectrum of single recurring events occurring in isolation. Neural activity is composed of many such events. We will thus now consider how multiple recurring events contribute to the voltage recorded by an electrode and how they collectively shape the Fourier spectrum of neural recordings. Let us illustrate how this may happen in a simple recording configuration.

As can be derived from Gauss’s law (the first of Maxwell’s equation), the voltage deflection caused by multiple sources at an electrode is the sum of the voltage induced by each source. Thus, the voltage induced by neural events simply sums up in the local field potential. The Fourier transform has the useful property that the spectrum of a sum of signals is the sum of their spectra. Multiple neural signals thus merely add their energy distribution to the spectrum of neural recordings. The voltage deflection induced by a single source is inversely proportional to its distance to the recording electrode. Sources that are small and relatively far away from the recording site, such as synaptic events, tend to be indistinguishable and sum into a background signal. We will thus first simulate this background signal.

In awake cortical recordings, background synaptic events often occur at a sustained high rate, a regime that is called desynchronized (Steriade and Deschenes, 1984; Destexhe and Paré, 1999; Chance, Abbott and Reyes, 2002; Destexhe and Contreras, 2006; Haider and McCormick, 2009; Petersen and Crochet, 2013). As synaptic responses tend to be slower, the spectrum has properties that are mostly shaped by the waveform of synaptic events (i.e., impulse response function, previous section). Synaptic events display a characteristic 1/f^2^ energy distribution (Figure 2B2). This energy distribution is inherited from the biophysical properties of neuronal membranes which tend to behave as passive resistor-capacitor (RC) filters (Koch, 1984). Here, we simply simulated the background LFP signal as white noise passing through an RC filter (Figure 3A1, Materials and Methods). The spectrum of that times series was computed as before (Figure 3A2).

Recordings of the local field potential will also often be impacted by action potentials originating from one or several neurons (i.e., units) located in the vicinity (within ∼30-50μm) of the recording electrode (Schmitzer-Torbert *et al*., 2005). The voltage deflection induced by these action potentials can sometimes have a significant impact on the spectrum. To illustrate this, we simulated neuronal firing by convolving a Poisson timing pulse train with a spike shaped Morlet wavelet. (Figure 3B1). As the timing of the train is Poissonian, the distribution of energy of the spectrum is purely inherited from the waveform of the Morlet spike (i.e., the impulse response function). The spectrum of the process showed a bell-shaped energy distribution with a peak at 75Hz (Figure 3B2).

The two time-series were then summed to simulate a common recording scenario where neuronal firing at the vicinity of the electrode adds to a background synaptic signal (Figure 3C1). The spectrum of the resulting times series was estimated as before (Figure 3C2). The spectrum of the sum of the two signal is nearly equivalent to the sum of their spectra. The Poisson spike train added to the spectrum of the background LFP an induced an upward inflection of the energy distribution peaking at 75Hz. This simulation illustrates how the contribution of multiple neuronal processes can added up in the Fourier Spectrum. These results also show that a relatively complex spectrum can arise from the simple addition of non-periodic processes.

### Identifying regular patterns in background synaptic activity

As illustrated in the example above, background synaptic activity has considerable influence on the shape of the LFP spectrum, especially at frequencies under ∼50Hz (Buzsáki, Anastassiou and Koch, 2012). We have also seen that under a desynchronized regime, the spectral energy of this background activity tends toward a 1/f^2^ distribution. However, background synaptic activity will at times include patterns of synchronized synaptic barrages (Buzsáki, Anastassiou and Koch, 2012; Buzsáki and Vöröslakos, 2023). These barrages tend to target specific subregions of the somato-dendritic compartment and thus, to induce return currents as neurons equilibrate their membrane potential. In oriented structures, such as the cortex and hippocampus, return currents add up and greatly amplify the voltage induce by barrages in the LFP (Buzsáki, Anastassiou and Koch, 2012). Thus, localized synaptic barrage often have a strong influence on the spectrum.

The temporal dynamics of synaptic barrages and their influence on the spectrum tend to vary from region to region and involve specific pathways. Characterizing these patterns and their influence on neural processing is a subject of active research (Girardeau *et al*., 2009; Cei *et al*., 2014; Veit *et al*., 2017, 2022; Uran *et al*., 2022; Buzsáki and Vöröslakos, 2023). Some well-characterized patterns of synaptic barrages show strong periodicity. These include thalamocortical spindles (Steriade and Deschenes, 1984; Timofeev, Contreras and Steriade, 1996; Steriade, 2006), the theta rhythm of the hippocampus (Vanderwolf, 1969; O’Keefe and Recce, 1993; Skaggs *et al*., 1996) or cortical activity during slow wave sleep (Steriade, Nuñez and Amzica, 1993; Contreras, Timofeev and Steriade, 1996). It is thus often natural to treat them as oscillations. However, we will illustrate how it is also useful and generally applicable to conceptualize these patterns as a train of recurring events such as those described in the previous sections. To illustrate this, we will use data that we have acquired in a recent study focusing on activity induced in the gamma range (30-80Hz) in the visual cortex of mice (Perrenoud *et al*., 2022).

Gamma activity arises in many brain regions under varying behavioral contingencies (Bosman *et al*., 2012; Cardin, 2016; Chen *et al*., 2017; Uran *et al*., 2022; Fernandez-Ruiz *et al*., 2023). In the mouse visual cortex, a specific increase of gamma activity is observed during locomotion (Niell and Stryker, 2010; Vinck *et al*., 2015). This activity comprises a narrow band component around 60Hz (Saleem *et al*., 2017; Shin *et al*., 2023) and a broad band component starting from ∼30 and expending to high frequencies (Figure 4B). In a recent study (Perrenoud *et al*., 2022), we took advantage of multichannel electrode arrays to show that this spectral increase can be tied to recurring events having a specific pattern of propagation across cortical layers (Figure 4A, C). Gamma events occurred more frequently during locomotion but also happened at a sustained rate during quiescence (Figure 4C). Decomposing gamma activity as an event train allows examination of how it entrains cortical neurons in relation with behavior with a high temporal resolution (Perrenoud *et al*., 2022) and provides some insight into how its spectral signature arises and contributes to the LFP.

Gamma events showed a complex inter-event distribution with a prominent mode around 17.8ms, showing that gamma events tend to occur in rapid sequences or bursts (Figure 4E1). Averaging the field potential around events shows a rhythmic pattern of activity (Figure 4D1), whose spectral energy distribution resembles that observed during locomotion (taking away the 1/f^2^ RC filtering induced by neural membrane, Figure 4D2). As gamma events shift phase across recording locations (Figure 4D1), it is difficult to precisely isolate the waveform of single events. However, we can now estimate how much of the shape of cortical gamma activity is determined by event timing. To do this we computed an idealized perfect pulse times series having a value of one at the time of events and zero everywhere else. The spectrum of that time series was computed as described above. We found that the characteristic broadband and narrow-band components of the distribution of energy of gamma activity can be derived solely by considering the events timing (Figure 4E2). Gamma activity in mouse V1, is thus a real-world example of how a complex spectrum can be shaped by the temporal distribution of recurring neural events.

## Discussion

Neural activity has highly complex dynamics (Buzsaki, 2006; Buzsáki, Anastassiou and Koch, 2012). The Fourier transform is a remarkably powerful tool allowing the detection of subtle changes in neural processes (Percival and Walden, 1993; Bruns, 2004). However, relating variation in the Fourier spectrum to actual neural processes is a complex problem. Thus, the interpretation of the spectrum of neural activity is rarely straightforward.

In the present study, we propose a simple yet generally applicable conceptual framework to guide the interpretation of the Fourier spectrum of neural recordings. We note that neural activity is made of recurring events such as synaptic currents, action potentials and, at larger scales, coordinated synaptic barrages within a region or from one region to another. Using simulations, we show that the spectrum of recurring neural events depends critically on two aspects: 1) the distribution of events in time and 2) the voltage deflection induced by single events. We detail how multiple recurring events simply sum up in the LFP and illustrate how this shapes the spectrum in a typical recording scenario. Finally, we illustrate how this framework can be used to describe the dynamics of gamma activity (30-80Hz) in real data obtained in the mouse visual cortex (Perrenoud *et al*., 2022).

Increased energy in the frequency spectrum within some frequency bands is often interpreted as the reflection of oscillatory neural activity. Accordingly, neural activity is frequently studied with concepts related to oscillations such as phase, amplitude, and coherence (Vinck *et al*., 2023). Some neural patterns do show strong periodicity (Buzsaki, 2006; Buzsáki and Vöröslakos, 2023), including slow wave sleep cortical activity (Steriade, Nuñez and Amzica, 1993; Contreras, Timofeev and Steriade, 1996; Sanchez-Vives and McCormick, 2000), theta rhythm in the hippocampus (O’Keefe and Recce, 1993; Skaggs *et al*., 1996) or thalamic spindles (Steriade, 2006; Niethard *et al*., 2018). Treating these patterns as oscillatory is thus a powerful way to quantify their properties (Bruns, 2004). However, these notions will not appropriately capture the properties of non-periodic signals (Donoghue *et al*., 2020). When then, might it be appropriate to treat neural patterns as oscillations?

Conceptualizing neural process as recurring events can a powerful way to address this question. Here, this framework allowed us to investigate how the spectrum is shaped by processes having arbitrary degrees of periodicity. Our work indicates that a tendency towards periodicity will indeed translate into increased power within a defined frequency band. However, we also provide simple examples of how band-specific power increase can arise from process that are entirely non-periodic. This indicates that some caution is warranted when concluding that peaks in the spectrum are the signature of oscillations or rhythmicity. One must indeed make sure that there is some energy in the spectrum within a specific frequency band. However, it is important that verification be performed in the time domain. A simple check for consistency in the amplitude and phase of the filtered signal can be an appropriate way to address these concerns.

What to do then when neural signals do not show strong signs of periodicity? Here, with the example of one of our recent studies (Perrenoud *et al*., 2022), we illustrate how treating of neural process as recurring events can also yield key insights into their properties (Figure 4). Thinking of neural patterns as recurring events is a straightforward way to link them to tangible neural processes such as synaptic responses, action potentials, or synaptic barrages. It also permits direct quantification of how the distribution of events in time might shape the spectrum. When the waveform of the voltage induced by single events can be estimated with precision (as for action potentials) it is potentially possible to infer and subtract the full contribution of a recurring event to the spectrum.

Identifying repeating events in the Local Field Potential and relating them to neural processes is not a simple problem. We and others have had success in identifying recurring events by taking advantage of multichannel recordings to resolve regularity in the propagation of neural activity in space (Perrenoud *et al*., 2022; Sibille *et al*., 2022). Recent progress in increasing the spatial resolution of multielectrode recordings may make this approach more feasible in the future (Jun *et al*., 2017; Steinmetz *et al*., 2021). Another important limitation of the framework introduced here is that, for the sake of clarity, we have only considered waveforms that are invariant in shape and amplitude. In a real-world scenario there might be variability in the intensity and shape of the waveform of recurring neural patterns over time. However, the simple conceptual framework introduced here may be a useful conceptual starting point to guide developments addressing these questions in the future.

## Data availability statement

The script and data used to generate the figure will be made available online (https://dryad).

## Ethics statement

All animal handling and experiments were performed according to the ethical guidelines of the Institutional Animal Care and Use Committee of the Yale University School of Medicine.

## Author contribution

QP and JAC designed the research, QP performed the simulations and wrote the manuscript, JAC acquired funding, supervised the project, and critically reviewed the manuscript.

## Funding

This work was supported by funding from the NIH (EY022951 and MH113852 to JAC, EY026878 to the Yale Vision Core), a McKnight Scholar Award (to JAC), an award from the Kavli Institute of Neuroscience (to JAC), and a BBRF Young Investigator Grant (to QP).

## Acknowledgement

The authors are grateful to Garrett Neske for helpful discussion and criticism.

---

**Supplementary Figure 1:**
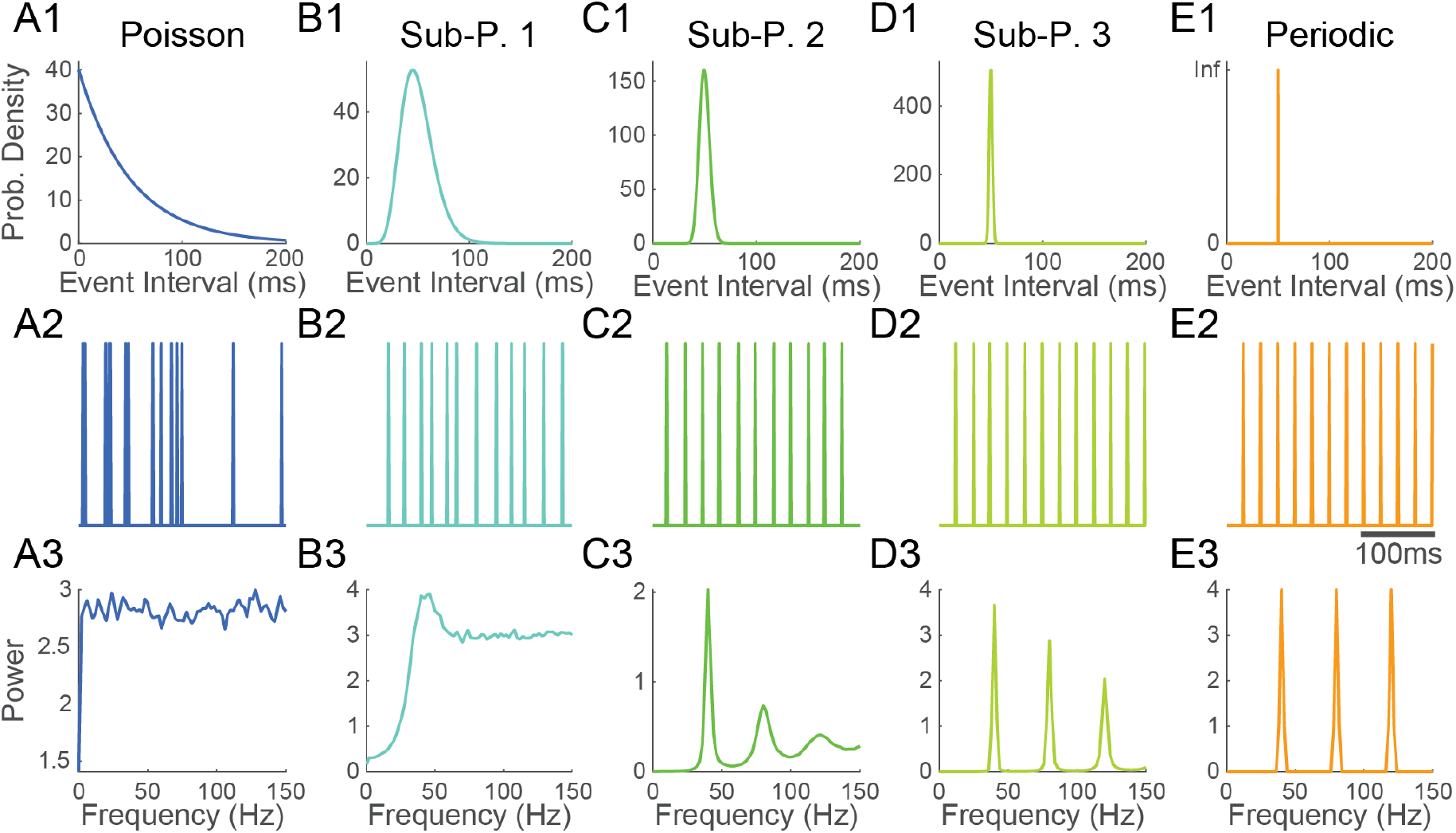
Transition from a Poisson to a periodic point process. **1)** Event interval distribution, **2)** excerpt of an event train and **3)** Fourier spectrum of **A)** a Poisson process, **B), C), D)** sub-Poissonian processes having diminishing variance in their inter-event interval distribution and **E)** a perfectly periodic point process. As the variance of the inter-event interval diminishes, secondary peak start to appear at the harmonics (i.e., integer multiples) of the mean frequency. All processes have a mean frequency of 40Hz.

**Supplementary Figure 2:**
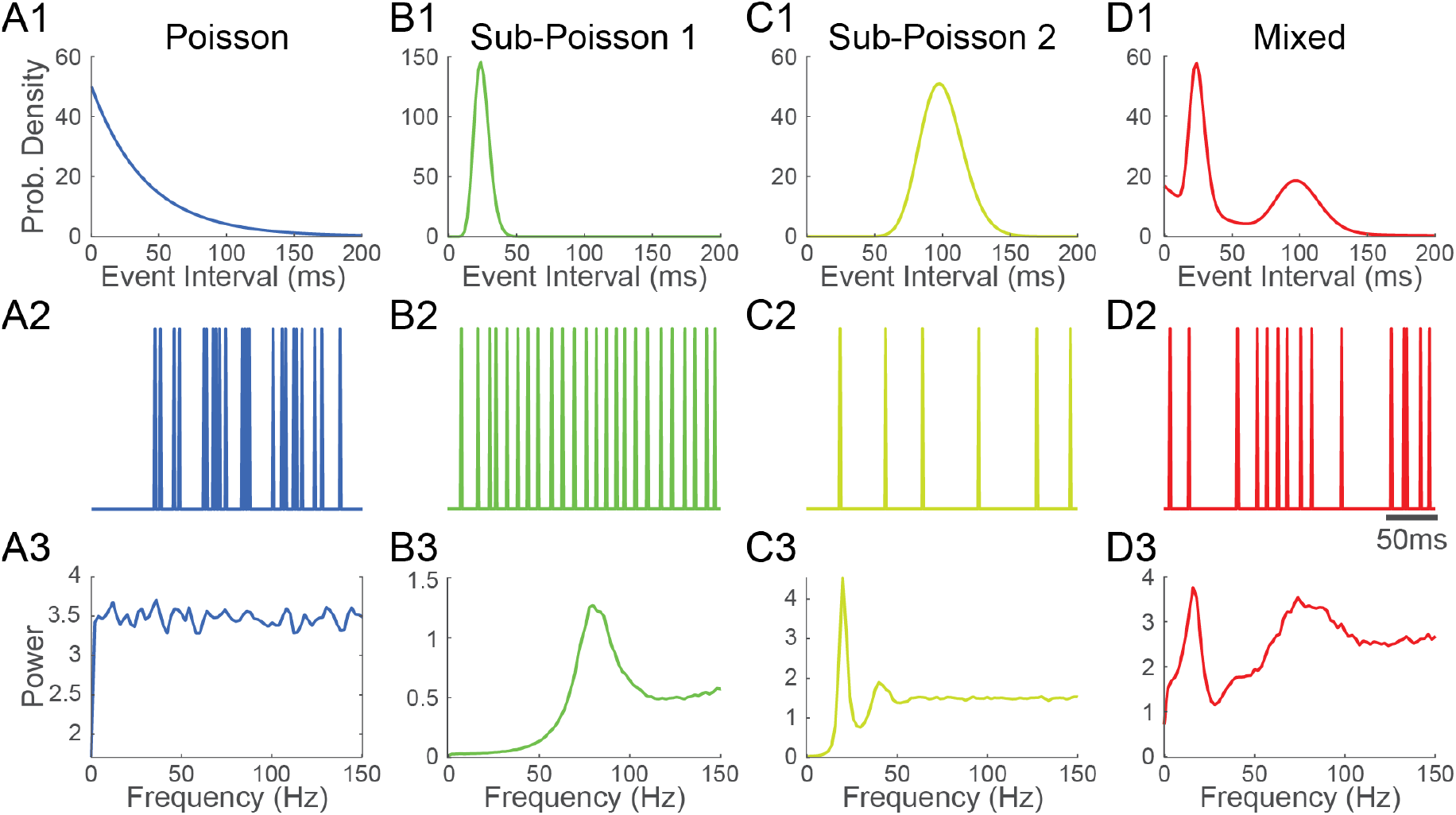
Peaks in the spectrum of a point process reflect peaks in the event interval distribution. **1)** Event interval distribution, **2)** excerpt of an event train and **3)** Fourier spectrum of **A)** a Poisson process (mean frequency 50Hz), **B), C)** two sub-Poissonian processes (mean frequency, 80Hz and 20Hz respectively) and **D)** a mixture of the processes in A, B and C. Peaks in the Fourier spectra of point processes indicate the presence of some preferred event interval.

**Supplementary Figure 3:**
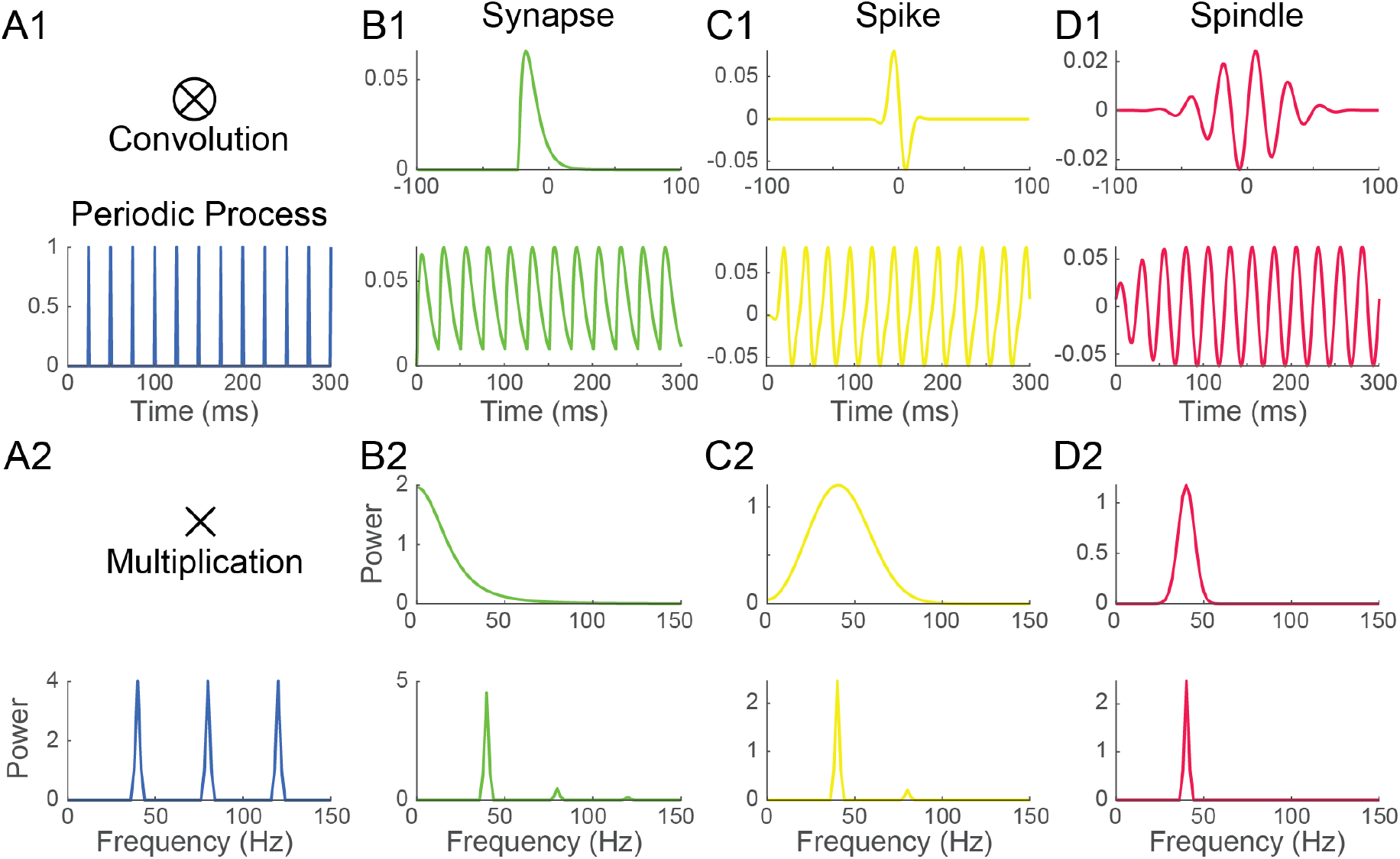
The Fourier spectrum of recurring event trains depends on the inter-event interval distribution and the waveform of single events. **1) A)** a perfectly periodic process was convolved with 3 waveforms (top) mimicking the shape of **B)** a synaptic event, **C)** a spike and **D)** a spindle, resulting in three distinct recurring event time series having the same event timing (bottom). **2)** Convolution in the time domain translates into a simple multiplication into the frequency domain. Thus, the Fourier spectrum of the recurring event time series in B1, C1 and D1 (bottom) is simply the product of the spectrum of their waveform (top) and the spectrum of the periodic pulse train (A2).

**Supplementary Figure 4:**
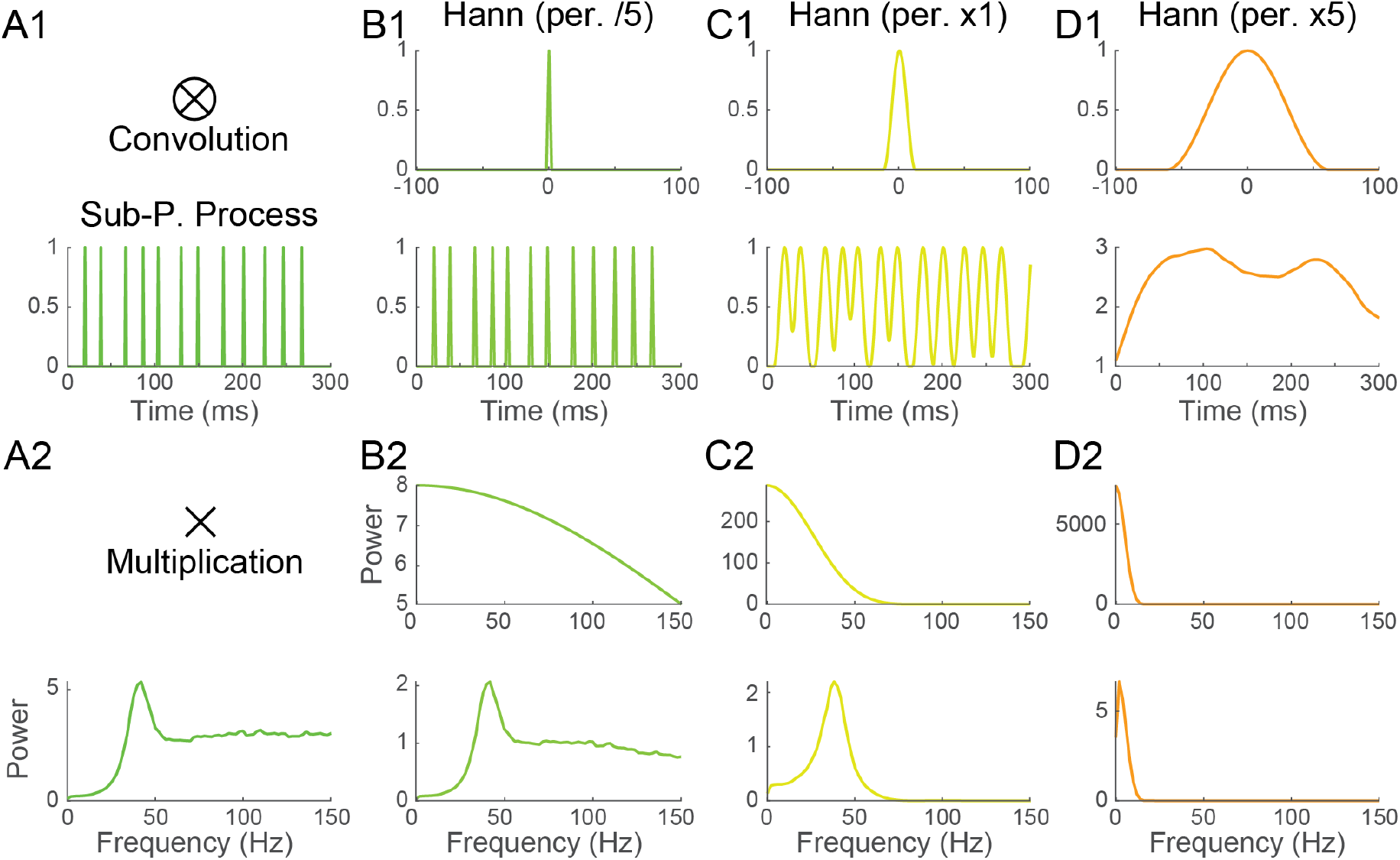
The width of the event waveform relative to the process rate influences how much each shapes the Fourier spectrum. **1) A)** a quasi-periodic sub-Poissonian process was convolved with 3 Hann waveform (top) having widths **B)** 5 time shorter, **C)** equal, and **D)** five time longer that the mean period of the process, resulting in three distinct recurring event time series having the same event timing (bottom). **2)** Convolution in the time domain translates into a simple multiplication into the frequency domain. Thus, the Fourier spectrum of the recurring event time series in B1, C1 and D1 (bottom) is simply the product of the spectrum of their impulse response functions (top) and the spectrum of the pulse train (A2). Shorter event waveforms make the spectrum look more like that of the event timing whereas longer waveforms tend to dominate the spectrum.

## Notes

### Competing Interest Statement

The authors have declared no competing interest.

